# SPECC1L-deficient palate mesenchyme cells show speed and directionality defect

**DOI:** 10.1101/854273

**Authors:** Jeremy P. Goering, Dona Greta Isai, Everett G. Hall, Nathan R. Wilson, Edina Kosa, Luke W. Wenger, Zaid Umar, Abdul Yousaf, Andras Czirok, Irfan Saadi

## Abstract

Clefts of the lip and/or palate (CL/P) are common anomalies that occur in 1/800 live births. Pathogenic *SPECC1L* variants identified in patients with rare atypical clefts and syndromic CL/P suggest the gene plays a primary role in face and palate development. We have generated *Specc1l* gene-trap (*Specc1l^cGT^*) and truncation (*Specc1l^ΔC510^*) alleles that cause embryonic or perinatal lethality, respectively. *Specc1l^cGT/ΔC510^* compound mutants show delayed and abnormal palatal shelf elevation at E14.5. By E15.5, the mutant shelves do elevate and fuse, however, the palatal rugae form abnormally. Palatogenesis requires extensive mesenchymal remodeling, especially during palatal shelf elevation. We posit that this remodeling involves collective movement of neural crest-derived palatal mesenchyme cells. Live time-lapse microscopy was performed to visualize in vitro wound-repair assays with wildtype and SPECC1L-deficient primary mouse embryonic palatal mesenchyme (MEPM) cells. SPECC1L-deficient MEPM cells consistently showed delayed closure in wound-repair assays. To evaluate which features of cellular movement were responsible, we performed automated particle image velocimetry (PIV) and manual cell tracking. The analyses revealed that both cell speed and directionality are disrupted in SPECC1L-deficient cells compared to controls. To determine if primary MEPM cells can move collectively, we assayed stream formation, which is a hallmark of collective movement. Indeed, MEPM cultures displayed correlated movement of neighboring cells. Importantly, correlation length was reduced in SPECC1L-deficient cultures, consistent with a role for SPECC1L in collective migration. Furthermore, we demonstrated that activation of the PI3K-AKT pathway with the 740Y-P small molecule can rescue the wound-closure delay in SPECC1L-deficient MEPM cells. Cell tracking analyses showed that this rescue was due to both increased speed and improved directionality. Altogether, our data showed a novel role for SPECC1L in guided movement through modulation of PI3K-AKT signaling.

## Introduction

Development of secondary palate involves coordinated growth and movement of palatal shelves in conjunction with surrounding craniofacial structures (Bush and Jiang 2012; Lan et al. 2015; Li et al. 2017a). In mice, the palatal shelves originate as a pair of vertical outgrowths from the maxillary processes at embryonic-day 11.5 (E11.5) and extend downward adjacent to the tongue until E13.5 (Bush and Jiang 2012). By E14.5, the palatal shelves elevate to position themselves horizontally above the tongue and extend toward the midline. By E15.5, the shelves have adhered and fused at the midline to form the secondary palate. Defects in palatal shelf outgrowth, elevation, or fusion can lead to cleft palate, one of the most common human birth defects (Bush and Jiang 2012; Mossey et al. 2009). Among these steps, elevation is the most poorly understood. Histological studies of palate elevation have shown that the mechanism differs along the anteroposterior axis (Bush and Jiang 2012; Jin et al. 2010; Yu and Ornitz 2011). The anterior palate exhibits a “flipping-up” motion where the distal ends of the shelves rise toward the midline. In contrast, the middle and posterior sections of the palate undergo more extensive remodeling, wherein the medial walls of the shelves extend horizontally as the distal ends of the shelves retract, creating a “bulge”(Jin et al. 2010; Walker and Fraser 1956; Yu and Ornitz 2011). This latter process can be referred to as mesenchymal remodeling (Jin et al. 2010). Interestingly, the “bulging” appears to occur first, indicating it may be the driving event during elevation (Yu and Ornitz 2011).

Several cellular mechanisms for mesenchymal remodeling during palate elevation have been proposed (Bush and Jiang 2012; Gritli-Linde 2008; Lan et al. 2015; Li et al. 2017b). Cell proliferation is required, but not sufficient, for elevation to occur (Jin et al. 2008; Kouskoura et al. 2013; Lan et al. 2016). Organ culture of palatal shelf explants have shown that palatal mesenchyme also has migratory properties, potentially guided by WNT5A and FGF10 chemotactic gradients (He et al. 2008). Thus, coordinated movement of the palate mesenchyme may contribute to palate remodeling, however, collective migration attributes have not been investigated in palate mesenchymal cells.

SPECC1L is a cytoskeletal protein that associates with both filamentous actin and microtubules (Saadi et al. 2011). Mutations in *SPECC1L* have been identified in multiple patients with craniofacial malformations, including cleft palate (Bhoj et al. 2018; Bhoj et al. 2015; Kruszka et al. 2015; Saadi et al. 2011). Mouse embryos homozygous for a null *Specc1l* gene-trap allele die between E9.5-E10.5 with open neural-folds and defects in cranial neural crest cell delamination (Wilson et al. 2016). Loss of SPECC1L also results in poor cell migration, increased filamentous actin staining, and abnormal staining of adherens junction markers β-catenin and E-cadherin (Saadi et al. 2011; Wilson et al. 2016). We have recently reported generation of a new gene-trap allele (*Specc1l^cGT^*), and a *Specc1l*-truncation allele lacking 510 C-terminal amino acids (*Specc1l^ΔC510^*). Homozygous mutants for *Specc1l^cGT^* are embryonic lethal, while those for *Specc1l^ΔC510^* are perinatal lethal (Hall EG In Revision). Crossing these alleles resulted in *Specc1l^cGT/ΔC510^* compound heterozygous mutant embryos that were also perinatal-lethal and exhibited a delay in palate elevation (Hall EG In Revision). As SPECC1L is broadly expressed in both palate epithelium and mesenchyme, we hypothesized that it may also play a role during mesenchymal remodeling.

In this study we used quantitative analyses of motility to show that primary mouse embryonic palatal mesenchyme (MEPM) cells exhibited stream formation, an attribute of collective migration, and showed that this behavior is impaired in *Specc1l*-mutant MEPM cells. We also performed wound-repair experiments using primary MEPM cells from *Specc1l^cGT/ΔC510^* mutant embryos to show defects in both cell speed and directionality. Importantly, we showed that pharmacological activation of the PI3K-AKT pathway, which is reduced in *Specc1l* mutants, is sufficient to rescue migration defects in these cells. Together, these data show a novel role for SPECC1L in cell migration through regulation of PI3K-AKT signaling pathway, as well as establish MEPM cells as a proxy model for mesenchymal remodeling during palate elevation.

## Materials and Methods

### MEPM isolation and cell culture

*Specc1l^ΔC510/+^* x *Specc1l^cGT/+^* mouse mating-pairs were placed together overnight, and the gestational stage was defined as E0.5 at noon on the day plug was identified. Females were euthanized at E13.5, and embryos were harvested in 1x PBS. Palatal shelves were dissected away from the oral cavity under aseptic conditions, placed in a 1.5mL tube with 0.5mL of 0.25% Trypsin (ThermoFisher, 25200056), and incubated at 37°C for 10 minutes. Occasional pipetting was performed to accelerate mechanical dissociation. The trypsinized MEPM cells were mixed with 5mL of high-glucose DMEM (HyClone, SH30243.01) containing 10% FBS (Corning, 35-010-CV) in a conical tube. The tube was centrifuged at 200rcf for 5 minutes to pellet the cells. MEPM cells were resuspended in fresh DMEM containing 10% FBS, plated on tissue-culture treated plastic dishes, and incubated at 37°C. Following expansion, cells were cryopreserved, and only passaged up to 3 times for use in experiments. All use of animal tissues followed an approved animal-use protocol.

### U2OS cell culture

Control and *SPECC1L*-kd U2OS cells were generated previously from U2OS osteosarcoma cells (ATCC HTB-96) as reported (Saadi et al. 2011). U2OS cells were cultured in DMEM (HyClone, SH30243.01) containing 10% FBS (Corning, 35-010-CV) as shown previously (Wilson et al. 2016).

### PI3K-AKT pathway activator treatment

For PI3K-AKT pathway small-molecule activation experiments, 100µg/mL 740Y-P (ApexBio, B5246) was added 24 hours prior to imaging. The activator containing medium was refreshed at the onset of imaging, and it remained throughout the duration of the experiment (48h). An equal proportion of water was added as vehicle to control wells.

### Live-cell imaging

Imaging was performed using the EVOS FL Auto Imaging System (ThermoFisher, AMAFD1000) with the EVOS Onstage Incubator (ThermoFisher, AMC1000). Cell cultures were kept in a humidified, 5% CO_2_ environment at 37°C over a period of 48-72 hours. Phase contrast images were collected at 4x or 10x magnification, every 10 or 20 minutes.

### Image processing

To detect cell-occupied area, a global threshold was applied to the local standard deviation of image brightness following the procedures outlined in Neufeld et al. (2017). The code is available at http://github.com/aczirok/cellconfluency. Cell motility was extracted using our particle image velocimetry (PIV) algorithm, with an initial window size of 50µm (Czirok et al. 2017; Zamir et al. 2005), resulting in a velocity field *v(x,t)* for each frame *t* and image location *x*. The average speed of cell motility was extracted from *v(x,t)* as a spatial average over the cell-occupied area.

### Wound-repair assay

For wound-repair assays, MEPM and U2OS cells were seeded at superconfluent densities of 1400/mm^2^ and 2300 cells/mm^2^ (respectively) into silicone inserts (Ibidi, 81176) in 35mm diameter tissue culture dishes. These silicone inserts provide 500µm wide cell-free gaps. Live-imaging started at the removal of the silicone insert, 48 hours after cell attachment (56 hours after seeding). For each wound, 4-5 non-overlapping fields were recorded every 10 minutes at 10x objective magnification. Image sequences were evaluated by compiling confluency data *A(t)* -- the size of cell-occupied area normalized to the field of view, as a function of time. The speed of wound-closure was established as *V = (w/2) (dA/dt)*, where *w* denotes the width of the field.

### Spontaneous collective motility assay

For the analysis of stream formation and collective motility, MEPM cells were seeded at low (30/mm^2^), medium (100/mm^2^), and high (300/mm^2^) densities into 3D-printed 6mm diameter rings on 35mm tissue culture dishes (Gulyas et al. 2018). Cultures were live-imaged every 20 minutes with a 4x phase contrast objective.

Local spatial correlations of cell movements were characterized by the average flow field that surrounds moving-cells as described previously (Czirok et al. 2013; Szabo et al. 2010). Briefly, a reference system was aligned to each vector *v(x,t)* and their “neighbors” registered in the appropriate bin (front, rear, etc.). The average velocity vector (*U*) in each bin is indicative of spatial correlation: the average of random vectors is close to zero, while it is non-zero when velocities share a common component. The calculated *U(x)* flow field was fitted with an exponential function *U(x)=a*exp(-x/x0)+U0*, where *x0* is the correlation length: the characteristic distance at which local velocity-velocity correlations disappear.

### Analysis of individual cell-trajectories

Individual cells were tracked manually on consecutive images (github/donnagreta/cm_track) yielding positions *P(i,t)* of cell *i* at time *t*. Motivated by a similar analysis performed by Biggs et al. (2014), trajectories were characterized by the total path-length 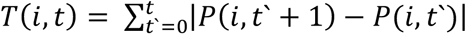, net displacement into the wound *D(i,t)=|X(i,t)-X(i,0)|*, where *X* is the projection of *P* in the direction perpendicular to the wound. Guidance efficiency was calculated as *D(i,t)/T(i,t)* for each cell *i* and timepoint *t*. Cultures were characterized by the population average of these single cell measures, evaluated at a suitable timepoint *t*.

## Results

### *Specc1l^cGT/ΔC510^* mutant embryos show abnormal palatogenesis

*Specc1l^cGT/ΔC510^* compound heterozygotes are perinatal lethal and show a delay in palate elevation at E14.5 (Hall EG In Revision). However, palatal shelves in most mutants recover and fuse by E15.5 (Fig.1) (Hall EG In Revision). Importantly, palate elevation in these mutants follows an abnormal sequence where posterior palate advances prior to middle and anterior palates, as in wildtype (WT) embryos (Fig. 1, b vs. e). Given the delayed and abnormal palate elevation in *Specc1l^cGT/ΔC510^* mutants, we hypothesized that (a) this delay is due to poor mesenchymal remodeling during elevation, and (b) coordinated movement of palatal mesenchyme cells is required for efficient remodeling. To determine the role of SPECC1L in the mesenchymal remodeling during palate elevation, we decided to perform *in vitro* motility assays using primary mouse embryonic palatal mesenchyme cells (MEPM) from WT and *Specc1l^cGT/ΔC510^* compound heterozygous embryos.

**Figure 1:**
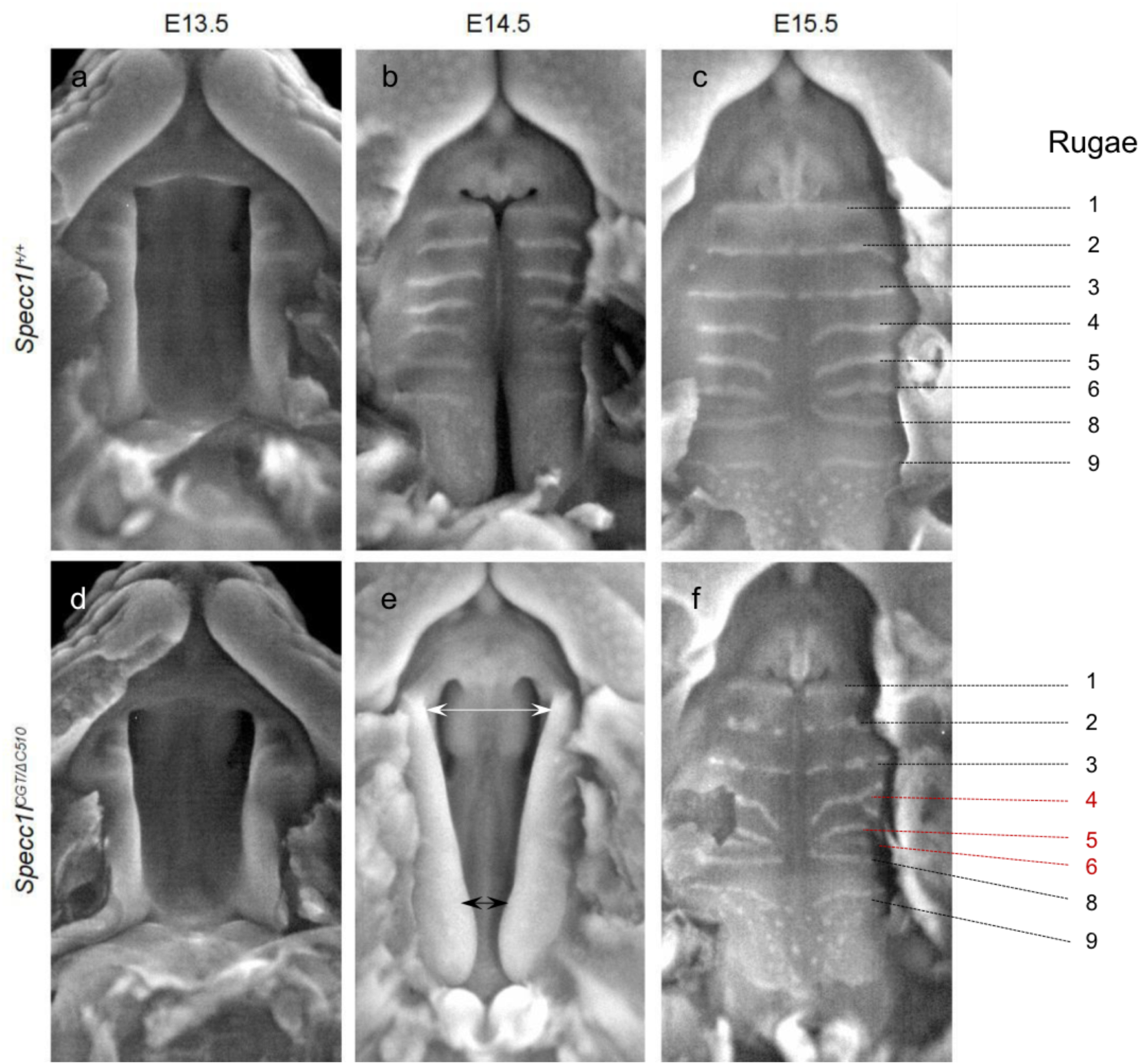
*Specc1l* deficiency results in abnormal palate closure and rugae formation. Compared to wildtype **(a-c)**, *Specc1l^cGT/ΔC510^* mutant embryos **(d-f)** exhibit abnormal palatal shelf elevation (b vs. e; arrows). By E14.5, normal palate elevation results in apposition of the anterior to middle regions of the palatal shelves (b). In contrast, at E14.5, mutant shelves show greater elevation posteriorly (e; black arrows) rather than in anterior and middle regions (e; white arrows). Although by E15.5, the mutant palatal shelves manage to elevate and adhere, they still show defects in rugae formation (f). These rugae patterning defects include both discontinuous and missing rugae (c vs. f), which persist until birth. Abnormal palatal shelf elevation sequence suggests a role for SPECC1L in mesenchymal remodeling during this process.

### *Specc1l*-deficient mouse embryonic palatal mesenchyme (MEPM) cells show migration defects during wound-closure

We previously reported that SPECC1L-deficient U2OS osteosarcoma cells showed poor migration in wound-repair assays (Saadi et al. 2011). To determine if MEPM cells from *Specc1l^cGT/ΔC510^* compound heterozygous embryos showed similar defects, we live-imaged wound-repair assays at superconfluent cell densities. As representative images of time-lapse recordings indicate (Fig. 2a), mutant MEPM cells take longer to close the wound than WT cells. This effect was quantified by image-analysis measures of confluency (Fig. 2b), from which a 33% reduction in the average speed of wound-fronts, from 6 µm/h to 4 µm/h, could be established (Fig. 2d). We used PIV analysis to quantify overall cell-motility within the entire microscopic field. The analysis indicated that MEPM cells remained motile even after closure of the wound (Fig. 2c), and the overall motile activity of *Specc1l^cGT/ΔC510^* mutant MEPM cells was reduced by 20%, from 4.4 µm/h to 3.5 µm/h (Fig. 2d).

**Figure 2.**
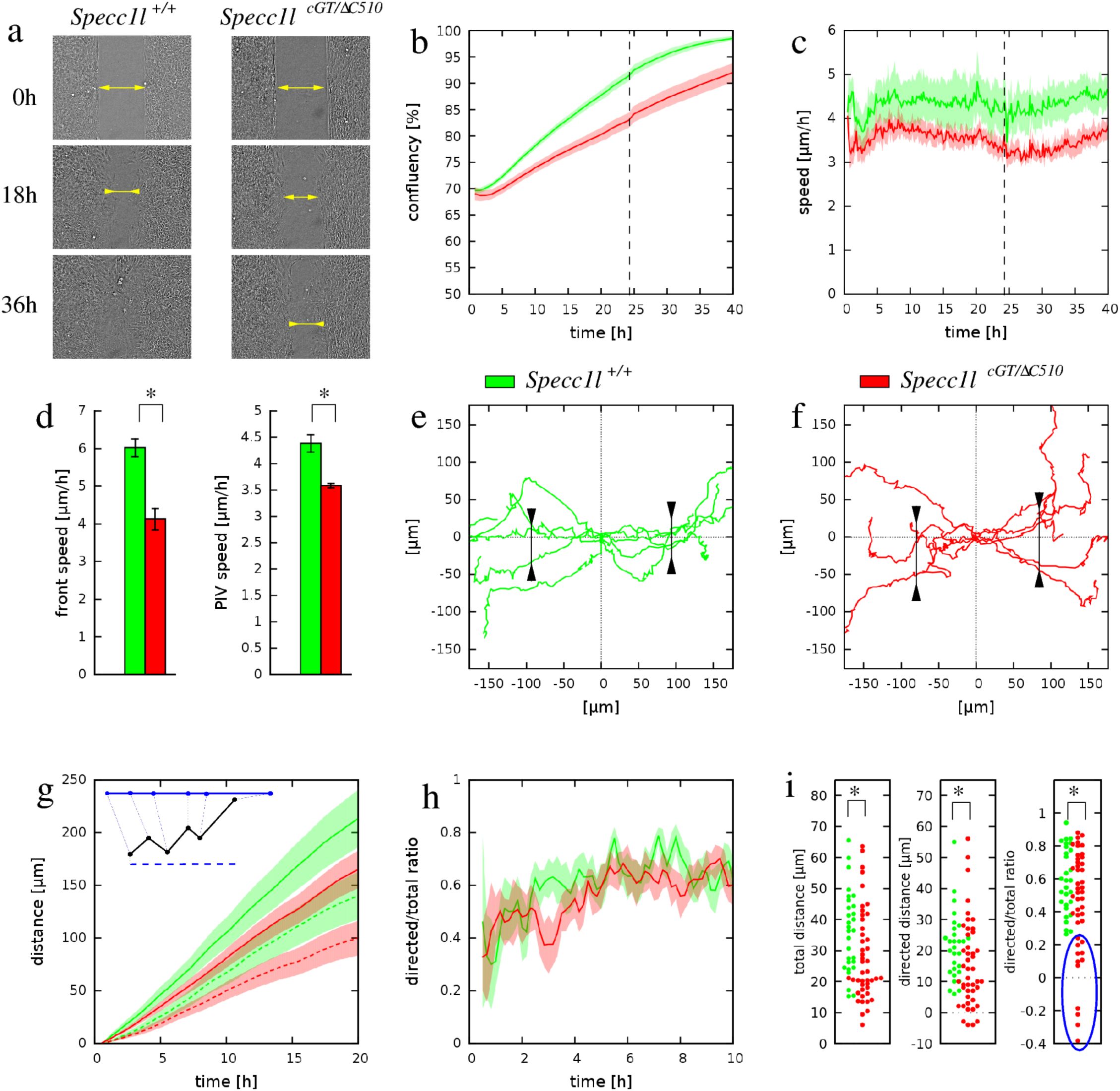
*Specc1l^cGT/ΔC510^* MEPM cells exhibit defects in directional migration. **a)** Phase contrast micrographs of a representative wound-repair assay performed with wild-type (WT) and *Specc1l*-mutant MEPM cells. While WT cells (left) show complete wound-closure by 36 hours (h), the mutant cells (right) require more time. The wound area is indicated by two-headed arrows. **b)** Automated confluence analysis of wound-repair experiments quantified the average percentage of wound-closure over time. Green and red colors indicate data obtained from WT and mutant cultures, respectively. By 24h (dashed vertical line), WT MEPM cells close more than 66% of the wound. Solid lines represent the average of 29 independent microscopic fields, recorded in 5 independent experiments. Shaded areas indicate SEM. **c)** Average cell motility speed, calculated by PIV analysis of the 29 wound-closure recordings analyzed in panel (b). For each time point velocity magnitudes within the entire cell-covered area were averaged. Despite approaching wound-closure after 24h (dashed line), cells remained motile in both WT and mutant MEPM cultures. **d)** Statistical analysis of data presented in panels (b-c) showed that the motility of mutant MEPM cells is significantly reduced in comparison to WT MEPM cells. Left: average speed of wound front, calculated within the 1h-15h time interval (p<6.7×10^−6^, Welch’s t-test). Right: average speed of cell movements within the entire recorded time period of 40h (p<2.2×10^−5^, Welch’s t-test). Error bars represent SEM. **e-f)** Representative paths of individual cells moving near the wound edge in WT (e) and mutant (f) cultures. Trajectories were shifted so that each originate at the origin. The WT cells exhibited a more directed movement to the open area compared to the mutant cells, as evidenced by the clustering of trajectories along the horizontal axis. Markers indicate the spread of trajectories: 90% of the trajectories pass through the drawn segments. **g)** Individual cell trajectories were statistically characterized by their mean total path lengths (solid lines) and net displacements directed into the wound area (dashed lines). The inset demonstrates these concepts with an example trajectory (black). Both measures were plotted as a function of time elapsed, and show a reduction of motile activity in *Specc1l*-mutant MEPM cells. Lines represent an average of 40 cell trajectories, shaded areas indicate SEM. **h)** The average ratio of the directed (net) displacement and total path length of the 40 cell trajectories, as a function of time. The typically lower values obtained for mutant MEPM cells indicate a guidance defect. **i)** Statistical analysis of individual cell trajectories using the measures of total displacement, directed displacement and their ratio. The bee swarm plots show these quantities for each tracked cell, evaluated at 10h culture time. The unguided population delineated by blue was present only in cultures of *Specc1l-*mutant MEPM cells. The significance of differences was established by pairwise Welsh’s t-tests, yielding p-values of 0.01, 0.05 and 0.03 for the total path length, directed displacement and ratio data sets, respectively.

To identify motility defects at the level of individual cells, 40 WT and 40 *Specc1l^cGT/ΔC510^* mutant MEPM cells were tracked manually as they moved into the wound. Cell trajectories of *Specc1l^cGT/ΔC510^* mutant MEPM cells (Fig. 2e,f) indicated a larger deviation perpendicular to the direction of wound-closure, hence a less-efficient guidance into the wound. Each trajectory was characterized at various timepoints during wound-closure by: (1) the length of total distance migrated (Fig. 2g), (2) the net displacement into the wound (Fig. 2g), and (3) the ratio of these two distances as a measure of guidance efficiency (Fig. 2h). While these quantities did not indicate a sudden change in cell behavior, the guidance efficiency of WT MEPM cells improved by ∼50% during the first 10h of the wound-closure process (Fig. 2h). Compared to these WT measures, the *Specc1l^cGT/ΔC510^* mutant MEPM cells exhibited a 25% reduction in total distance migrated, and a larger 30% reduction in the net displacements into the wound (Fig. 2g). The larger reduction in directed motility is associated with consistently poor guidance efficiency of mutant MEPM cells (Fig. 2h). Distribution of the three motility measures within the tracked-cell population is quite broad (Fig. 2i). The most conspicuous difference between WT and *Specc1l^cGT/ΔC510^* mutant cultures is the presence of an unguided cell population within the latter (Fig. 2i, blue circle). Thus, population-level measures as well as analysis of individual trajectories indicate a substantial role of SPECC1L in both promoting MEPM migration and responding to guidance cues.

### MEPM cells display attributes of collective movement

Since our analysis of wound-repair live-imaging studies indicated that *Specc1l^cGT/ΔC510^* mutant cells had defects in speed and guidance, we asked a more fundamental question: do MEPM cells display attributes of collective movement? To test this, we recorded WT and *Specc1l^cGT/ΔC510^* mutant MEPM cells, seeded at various cell densities, in a simple two-dimensional tissue-culture environment (Fig. 3a-c). Interestingly, at high density, MEPM cells created ordered and long-range oriented domains reaching up to 1mm (Fig 3c).

**Figure 3.**
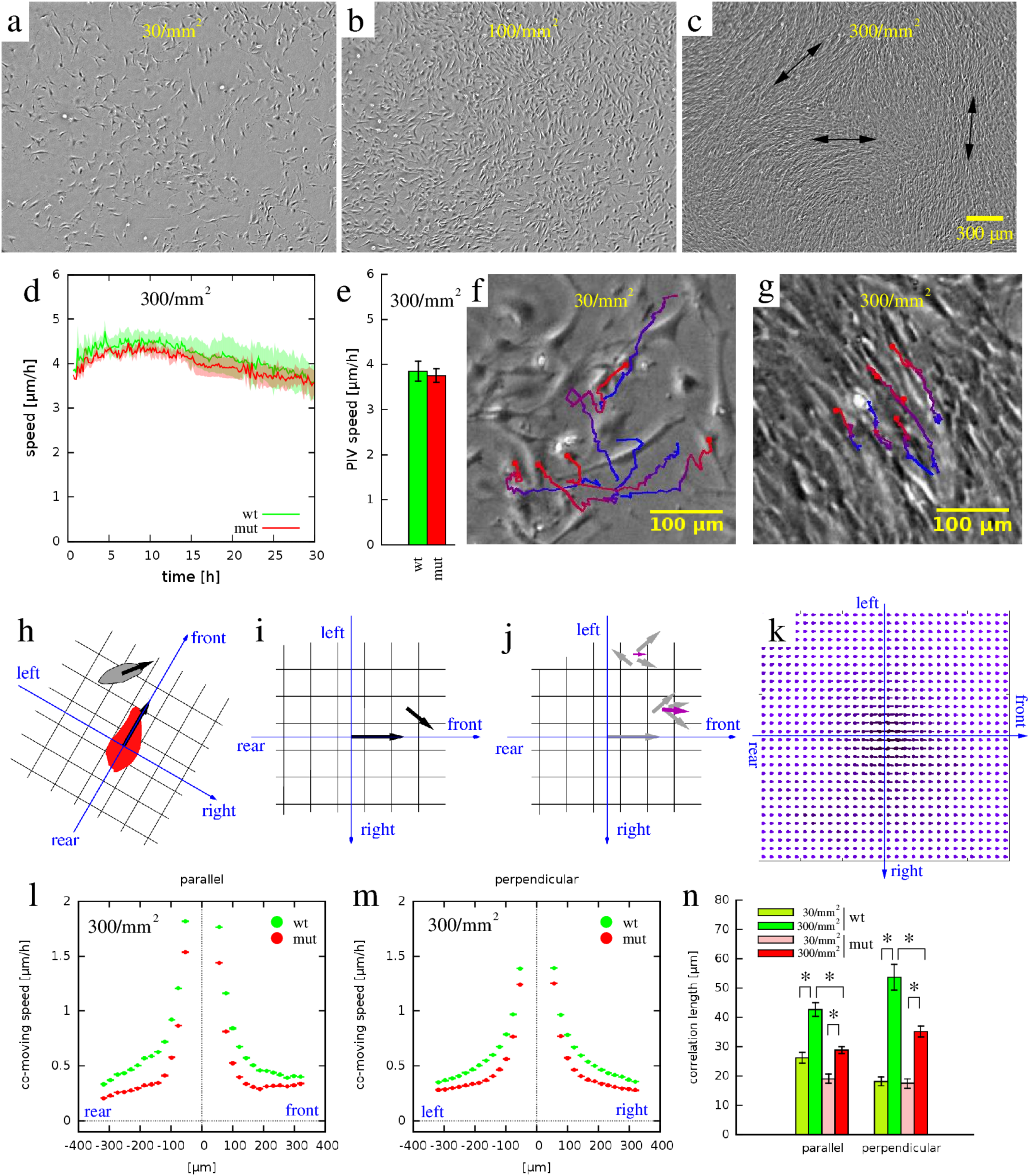
*Specc1l^cGT/ΔC510^* mutant MEPM cells exhibit a collective migration defect in 2D random motility assays. **a-c)** Representative phase contrast micrographs of wildtype (WT) MEPM cells at low (30/mm^2^), medium (100/mm^2^), and high (300/mm^2^) plating densities. When cultured at high cell densities, MEPM cells exhibited a local directional alignment of cells (arrows). **d)** The average cell motility speed of high density MEPM cultures demonstrated sustained cell motility despite the high cell density. Velocities were calculated by PIV analysis and averaged over the entire field of view. Green and red colors indicate WT and *Specc1l*-mutant cells, respectively. Solid lines are averages of 4 independent fields, shaded areas indicate SEM. **e)** The average speed of both WT and mutant cells were calculated for the entire recorded time period. Error bars indicate SEM, the difference is not significant. **f-g)** Tracks of individual cells in low (f) and high density (g) cultures. The random and uncorrelated motility at low cell densities (f) became highly aligned at high densities evidenced by the parallel trajectories (g). Trajectories are plotted for a time period of 8h, with red and blue colors indicating earlier and later segments, respectively. **h-k)** To characterize spatial correlations in cell motility, we determined the average co-moving velocity at various locations relative to an average motile cell. h) For each moving cell (red) we determined the relative position and velocity of every other cell (gray) within the vicinity. i) The velocities were assigned to bins of a coordinate system, co-aligned with the moving cell. j) By repeating this procedure for multiple cells and multiple time points, each bin contained multiple velocity vectors (gray). The average velocity vector in each bin (magenta) was indicative of the spatial correlation: the average of random vectors was close to zero, while it was non-zero when velocities shared a common component. k) The average velocity map thus characterized the typical cell velocities at various locations relative to a moving cell. We sampled this field along the front-rear axis and also along the left-right axis (l, m). **l,m)** MEPM cells in high density cultures tended to move locally in the same direction, indicated by positive average co-moving speed values -- both along the front-rear (l) and lateral (m) axes. Data were averaged from 4 independent fields, both for WT (green) and *Specc1l*-mutant (red) cells. The co-movement of *Specc1l* -mutant cells, however, diminished at a closer distance than that of WT cells. **n)** Average correlation lengths, both parallel with and perpendicular to the direction of cell motion, were established by fitting an exponential function on the profiles shown in panels (l, m). High-density cultures (green and red) showed a significantly increased correlation length compared to low-density cultures (lime and pink). The increase, however, was approximately twice as large in WT cells (lime vs green) than in *Specc1l*-mutant cells (pink vs red). Significance values of the differences were established by Welch t-tests, and are summarized in Supplemental Table 1.

Spontaneous cell motility was quantified by PIV analysis, revealing a sustained motile behavior despite the high cell density (Fig 3d). The average motile speed was not significantly different between WT and mutant cultures (Fig 3e, ∼4 µm/h), in contrast to PIV cell speeds observed in wound-repair experiments (Fig 2d).

Trajectories of individually tracked cells revealed a cell-density-dependent ordering: cell trajectories of adjacent cells in low-density cultures appeared uncorrelated (Fig 3f), while those in high-density cultures were largely parallel (Fig 3g). To quantify this apparent collective motion of MEPM cells, we adopted a spatial cross-correlation measure (Czirok et al. 2013; Szabo et al. 2010), which determined the average co-movement speed of cells at various locations (e.g., front, rear, left, right) relative to a moving cell (Fig. 3h-k). Correlation between cell movements was higher when cells were in close vicinity and gradually tapered off as distance increased beyond 300µm (Fig. 3l,m). This effect was quantified by fitting the co-movement speed profiles with an exponential function, yielding the correlation length as a fitted parameter. Correlation length values indicated a 33% reduction in both the parallel and perpendicular correlation lengths of *Specc1l^cGT/ΔC510^* mutant MEPM cells when compared with WT MEPM cells. This reduction in correlated movement at high cell-densities (Fig. 3n) suggests an impaired ability of mutant cells to form streams.

### Upregulation of PI3K-AKT signaling rescues *Specc1l*-deficient migration defects

We previously reported that loss of SPECC1L results in decreased PI3K-AKT signaling along with defects in cell-adhesion and cell-shape (Wilson et al. 2016). Pharmacological activation of the pathway was sufficient to rescue these phenotypes (Wilson et al. 2016). To test if PI3K-AKT pathway activation could also rescue wound-closure defects in mutant MEPM cells, we repeated the wound-closure experiments reported in Fig.2 in the presence and absence of 740Y-P (100 µg/ml), an activator of PI3K (Fig. 4). We found that activation of PI3K-AKT pathway did indeed rescue the wound-closure defect in *Specc1l^cGT/ΔC510^* mutant MEPM cultures (Fig. 4a). Specifically, 740Y-P treatment increased the rate of wound-closure in both WT and *Specc1l^cGT/ΔC510^* mutant MEPM cultures by almost two-fold (Fig. 4c), even beyond the rate observed in untreated WT MEPM cultures. Interestingly, PIV analysis of 740Y-P treatment did not show an increase in overall WT cell motility, and only a 20% increase in *Specc1l^cGT/ΔC510^* mutant cell motility (Fig. 4b,c). Therefore, we hypothesized that improved cell-guidance into the wound contributed to the improved wound-closure in mutant cultures treated with 740Y-P.

**Figure 4.**
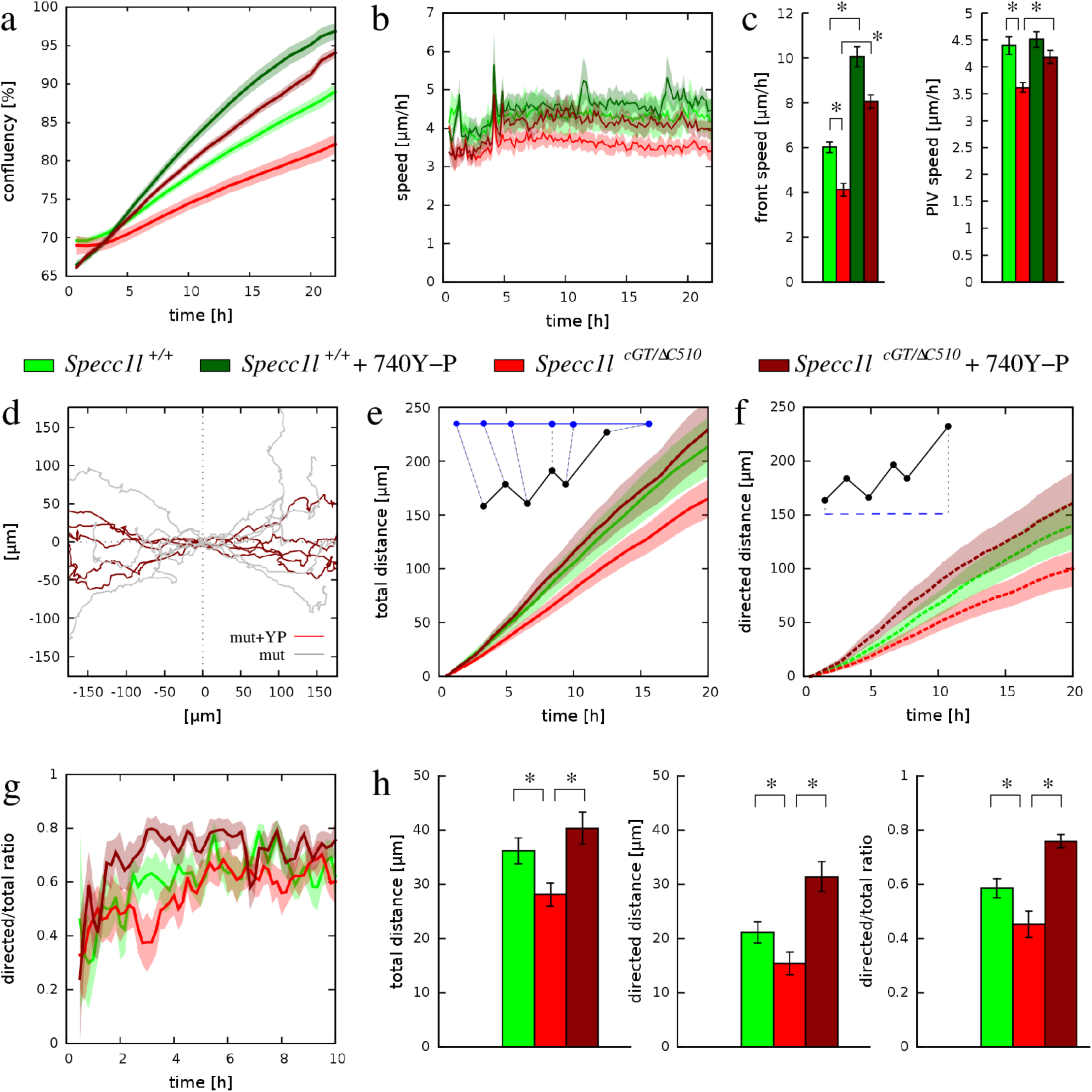
Upregulation of PI3K-AKT signaling rescues both speed and directionality of *Specc1l^cGT/ΔC510^* mutant cells in wound-closure assays. **a)** Confluency analysis of wound-closure recordings indicated a substantially increased speed of the process as a response of 100 µg/ml 740Y-P, a PI3K activator. Green and red colors indicate data obtained from wild-type (WT) and *Specc1l-*mutant cultures, while lighter and darker colors indicate vehicle-treated and 740Y-P treated cultures, respectively. Solid lines represent the average of 13 independent microscopic fields, recorded in 4 independent experiments. Shaded areas indicate SEM. **b)** Average cell motility speed, calculated by PIV analysis of the wound-closure recordings analyzed in panel (a). The activator 740Y-P induced an increased motile activity, persistent for at least a day, in *Specc1l*-mutant cells. **c)** Statistical analysis of data presented in panels (a,b) indicated that the motility defect of mutant MEPM cells was rescued by the PI3K-AKT activator 740Y-P. Left: average speed of front propagation, calculated within the 1h-15h time interval. The speed-up in wound-closure is significant (p<5×10^−7^ for WT and p<3×10^−10^ for mutant, Welch’s t-test). Right: average speed of cell movements within the entire recorded time period of 24h. The increase of speed was significant in *Specc1l*-mutant cells (p<4×10^−8^, Welch’s t-test). Error bars represent SEM. **d)** Trajectories of individual, 740Y-P treated *Specc1l*-mutant MEPM cells (dark red) was compared with that of untreated MEPM cells (gray). The PI3K-AKT activator enhanced the guidance of cell movements into the wound area. **e-h)** Statistical analysis of individual cell trajectories. Total path length (e), net directed displacement into the wound (f) and their ratio (g) indicated a sustained improvement in the performance of *Specc1l*-mutant MEPM cells. h) Bar charts indicating the cell trajectory measures evaluated at 10h after the onset of migration. The significance of rescue was established by pairwise Welsh’s t-tests, yielding p-values of 2×10^−3^, 2×10^−5^, and 4×10^−7^ for the total path length, net directed displacement and ratio data sets, respectively. The significance values for the WT-mutant comparison were reported in Fig 2.

To determine if cell-guidance was improved in the rescue experiments, we compared the trajectories of individual MEPM cells from treated and untreated cultures (Fig. 4d). The dispersion of *Specc1l^cGT/ΔC510^* mutant cells perpendicular to the direction of wound-closure was visibly reduced by PI3K activation, indicating improved guidance into the wound. Statistical evaluation of individual cell-trajectories (Fig. 4e-h) revealed a 40% increase in the total distance migrated (Fig. 4e,h), a two-fold increase in the net displacement directed into the wound (Fig. 4f,h), and a 66% increase in guidance efficiency (Fig. 4g,h) in 740Y-P treated *Specc1l^cGT/ΔC510^* mutant MEPM cells. Taken together, the data indicate that PI3K activation can rescue SPECC1L deficiency-related motility defects by improving both cell motility speed and guidance efficiency.

### SPECC1L-dependent collective migration in other cell types

SPECC1L deficiency has been previously shown to result in defects in cell migration in cultured U2OS cells (Saadi et al. 2011; Wilson et al. 2016). To support a primary role of SPECC1L in collective cell migration, we analyzed time-lapse imaging of control and *SPECC1L*-kd U2OS cells in wound-repair assays, similarly to the MEPM data. We were able to show again in U2OS cells that SPECC1L deficiency resulted in slower wound-closure coupled with poor stream-formation and directional movement (Suppl. Fig.1). We also confirmed that activation of the PI3K-AKT pathway rescued the cell speed and guidance defects in *SPECC1L*-kd U2OS cells (Suppl. Fig.2). These findings point to a common role of SPECC1L in the regulation of cell motility and guidance.

## Discussion

Studies of palatogenesis have demonstrated that remodeling from vertical to horizontal direction is a primary feature of the elevation process (Chiquet et al. 2016; Li et al. 2017a; Yu and Ornitz 2011). This remodeling is likely initiated by the medial edge epithelium of the palatal shelf, but driven by the palate mesenchyme. While primary MEPM cells have been used for testing the effect of various inhibitory compounds, growth factors, and gene-expression changes (Fantauzzo and Soriano 2014; Gao et al. 2019; Iyyanar and Nazarali 2017; Jiang et al. 2017; Liu et al. 2014; Vasudevan and Soriano 2014), only a few studies have investigated the migration of MEPM cells (Fantauzzo and Soriano 2017; Gao et al. 2019; He and Soriano 2013), and none, to our knowledge, have examined if they can migrate collectively. We identify the presence of stream formation in MEPM cells, a behavior generally considered to be a feature of collective movement (Czirok et al. 2013; Scarpa and Mayor 2016; Szabo et al. 2010). This process requires active cell-cell communication involving cell-adhesion molecules, as well as dynamic reorganization of the cytoskeleton to direct movement in response to signals from neighboring cells (Czirok et al. 2013; Tang and Gerlach 2017; Theveneau and Mayor 2013). The more pronounced correlation in movement seen in high-density MEPM cultures is consistent with this contact-dependent mechanism. Thus, our data indicate that cultured MEPM cells have an innate ability to move collectively.

We also show that SPECC1L deficiency impaired the ability of palatal mesenchyme cells to move collectively – a result that we posit partially explains the observed delayed and abnormal palate elevation in *Specc1l^cGT/ΔC510^* mutants (Fig.1) (Hall EG In Revision). The defect in collective migration was also validated in U2OS cells, an independent cell line, which strongly supports a primary role for SPECC1L in collective movement.

We previously showed that SPECC1L deficiency leads to reduced PI3K-AKT signaling, and that PI3K-AKT activation was able to rescue cell-shape and cell-adhesion phenotypes in *SPECC1L*-kd U2OS cells (Wilson et al. 2016). Our analyses now show that activation of PI3K-AKT signaling rescued both cell speed and directionality defects during wound-closure of *Specc1l^cGT/ΔC510^* mutant MEPM cells, as well as in *SPECC1L*-kd U2OS cells. Thus, our results are consistent with a pro-migratory role for AKT (Xue and Hemmings 2013) and suggest that SPECC1L regulates cell migration through PI3K-AKT signaling.

This study represents the first depiction of collective movement characteristics in cultured MEPM cells, and supports the hypothesis that palate elevation involves collective movement of the mesenchyme. In addition to our *Specc1l* mutants, there remain several mouse mutants with elevation defects for which the underlying pathogenic mechanisms have not been fully elucidated (Bush and Jiang 2012; Funato et al. 2015; Gritli-Linde 2008; Li et al. 2017b). Future analysis of these mutants may benefit from using primary MEPM cell migration as a proxy for mesenchymal remodeling during palatal shelf elevation.

## Acknowledgements

This project was supported in part by the National Institutes of Health grants DE026172 (I.S.), GM102801 (A.C.), and F31DE027284 (E.H.). I.S. was also supported in part by the Center of Biomedical Research Excellence (COBRE) grant (National Institute of General Medical Sciences P20 GM104936), Kansas IDeA Network for Biomedical Research Excellence grant (National Institute of General Medical Sciences P20 GM103418), and Kansas Intellectual and Developmental Disabilities Research Center (KIDDRC) grant (U54 Eunice Kennedy Shriver National Institute of Child Health and Human Development, HD 090216). The Confocal Imaging Facility, the Integrated Imaging Core, and the Transgenic and Gene Targeting Institutional Facility at the University of Kansas Medical Center are supported, in part, by NIH/NIGMS COBRE grant P30GM122731 and by NIH/NICHD KIDDRC grant U54HD090216.

## Author contributions

JPG, DGI, EGH, NRW, AC, and IS conceived and designed the experiments. JPG, EGH, EK, NRW, and LWW performed the experiments. DGI and AC performed the time-lapse imaging analysis. JPG, ZU, and AY performed the manual cell-tracking. EGH, AC and IS wrote the paper. JPG, DGI, NRW and LWW edited the manuscript. All authors reviewed the manuscript.

## References

Bhoj EJ, Haye D, Toutain A, Bonneau D, Nielsen IK, Lund IB, Bogaard P, Leenskjold S, Karaer K, Wild KT et al. 2018. Phenotypic spectrum associated with specc1l pathogenic variants: New families and critical review of the nosology of teebi, opitz gbbb, and baraitser-winter syndromes. Eur J Med Genet.

Bhoj EJ, Li D, Harr MH, Tian L, Wang T, Zhao Y, Qiu H, Kim C, Hoffman JD, Hakonarson H et al. 2015. Expanding the specc1l mutation phenotypic spectrum to include teebi hypertelorism syndrome. Am J Med Genet A. 167A(11):2497–2502.

Biggs LC, Naridze RL, DeMali KA, Lusche DF, Kuhl S, Soll DR, Schutte BC, Dunnwald M. 2014. Interferon regulatory factor 6 regulates keratinocyte migration. J Cell Sci. 127(Pt 13):2840–2848.

Bush JO, Jiang R. 2012. Palatogenesis: Morphogenetic and molecular mechanisms of secondary palate development. Development. 139(2):231–243.

Chiquet M, Blumer S, Angelini M, Mitsiadis TA, Katsaros C. 2016. Mesenchymal remodeling during palatal shelf elevation revealed by extracellular matrix and f-actin expression patterns. Front Physiol. 7:392.

Czirok A, Isai DG, Kosa E, Rajasingh S, Kinsey W, Neufeld Z, Rajasingh J. 2017. Optical-flow based non-invasive analysis of cardiomyocyte contractility. Sci Rep. 7(1):10404.

Czirok A, Varga K, Mehes E, Szabo A. 2013. Collective cell streams in epithelial monolayers depend on cell adhesion. New J Phys. 15:75006.

Fantauzzo KA, Soriano P. 2014. Pi3k-mediated pdgfralpha signaling regulates survival and proliferation in skeletal development through p53-dependent intracellular pathways. Genes Dev. 28(9):1005–1017.

Fantauzzo KA, Soriano P. 2017. Generation of an immortalized mouse embryonic palatal mesenchyme cell line. PLoS One. 12(6):e0179078.

Funato N, Nakamura M, Yanagisawa H. 2015. Molecular basis of cleft palates in mice. World J Biol Chem. 6(3):121–138.

Gao L, Xu J, Li X, Wang T, Wu W, Cao J. 2019. 2,3,7,8-tetrachlorodibenzo-p-dioxin and tgfbeta3-mediated mouse embryonic palatal mesenchymal cells. Dose Response. 17(1):1559325818786822.

Gritli-Linde A. 2008. The etiopathogenesis of cleft lip and cleft palate: Usefulness and caveats of mouse models. Curr Top Dev Biol. 84:37–138.

Gulyas M, Csiszer M, Mehes E, Czirok A. 2018. Software tools for cell culture-related 3d printed structures. PLoS One. 13(9):e0203203.

Hall EG WL, Wilson NR, Undurty-Akella S, Standley J, Augustine-Akpan EA, Kousa YA, Acevedo DS, Goering JP, Pitstick L, Natsume N, Paroya SM, Busch TD, Ito M, Mori A, Imura H, Schultz-Rogers LE, Klee EW, Babovic-Vuksanovic D, Kroc SA, Adeyemo WL, Eshete MA, Bjork BC, Suzuki S, Murray JC, Schutte BC, Butali A, Saadi I. In Revision. Specc1l regulates palate development downstream of irf6.

He F, Soriano P. 2013. A critical role for pdgfralpha signaling in medial nasal process development. PLoS Genet. 9(9):e1003851.

He F, Xiong W, Yu X, Espinoza-Lewis R, Liu C, Gu S, Nishita M, Suzuki K, Yamada G, Minami Y et al. 2008. Wnt5a regulates directional cell migration and cell proliferation via ror2-mediated noncanonical pathway in mammalian palate development. Development. 135(23):3871–3879.

Iyyanar PPR, Nazarali AJ. 2017. Hoxa2 inhibits bone morphogenetic protein signaling during osteogenic differentiation of the palatal mesenchyme. Front Physiol. 8:929.

Jiang Z, Pan L, Chen X, Chen Z, Xu D. 2017. Wnt6 influences the viability of mouse embryonic palatal mesenchymal cells via the beta-catenin pathway. Exp Ther Med. 14(6):5339–5344.

Jin JZ, Li Q, Higashi Y, Darling DS, Ding J. 2008. Analysis of zfhx1a mutant mice reveals palatal shelf contact-independent medial edge epithelial differentiation during palate fusion. Cell Tissue Res. 333(1):29–38.

Jin JZ, Tan M, Warner DR, Darling DS, Higashi Y, Gridley T, Ding J. 2010. Mesenchymal cell remodeling during mouse secondary palate reorientation. Dev Dyn. 239(7):2110–2117.

Kouskoura T, Kozlova A, Alexiou M, Blumer S, Zouvelou V, Katsaros C, Chiquet M, Mitsiadis TA, Graf D. 2013. The etiology of cleft palate formation in bmp7-deficient mice. PLoS One. 8(3):e59463.

Kruszka P, Li D, Harr MH, Wilson NR, Swarr D, McCormick EM, Chiavacci RM, Li M, Martinez AF, Hart RA et al. 2015. Mutations in specc1l, encoding sperm antigen with calponin homology and coiled-coil domains 1-like, are found in some cases of autosomal dominant opitz g/bbb syndrome. J Med Genet. 52(2):104–110.

Lan Y, Xu J, Jiang R. 2015. Cellular and molecular mechanisms of palatogenesis. Curr Top Dev Biol. 115:59–84.

Lan Y, Zhang N, Liu H, Xu J, Jiang R. 2016. Golgb1 regulates protein glycosylation and is crucial for mammalian palate development. Development. 143(13):2344–2355.

Li C, Lan Y, Jiang R. 2017a. Molecular and cellular mechanisms of palate development. J Dent Res. 96(11):1184–1191.

Li C, Lan Y, Krumlauf R, Jiang R. 2017b. Modulating wnt signaling rescues palate morphogenesis in pax9 mutant mice. J Dent Res. 96(11):1273–1281.

Liu X, Zhang H, Gao L, Yin Y, Pan X, Li Z, Li N, Li H, Yu Z. 2014. Negative interplay of retinoic acid and tgf-beta signaling mediated by tg-interacting factor to modulate mouse embryonic palate mesenchymal-cell proliferation. Birth Defects Res B Dev Reprod Toxicol. 101(6):403–409.

Mossey PA, Little J, Munger RG, Dixon MJ, Shaw WC. 2009. Cleft lip and palate. Lancet. 374(9703):1773–1785.

Neufeld Z, von Witt W, Lakatos D, Wang J, Hegedus B, Czirok A. 2017. The role of allee effect in modelling post resection recurrence of glioblastoma. PLoS Comput Biol. 13(11):e1005818.

Saadi I, Alkuraya FS, Gisselbrecht SS, Goessling W, Cavallesco R, Turbe-Doan A, Petrin AL, Harris J, Siddiqui U, Grix AW, Jr. et al. 2011. Deficiency of the cytoskeletal protein specc1l leads to oblique facial clefting. Am J Hum Genet. 89(1):44–55.

Scarpa E, Mayor R. 2016. Collective cell migration in development. J Cell Biol. 212(2):143–155.

Szabo A, Unnep R, Mehes E, Twal WO, Argraves WS, Cao Y, Czirok A. 2010. Collective cell motion in endothelial monolayers. Phys Biol. 7(4):046007.

Tang DD, Gerlach BD. 2017. The roles and regulation of the actin cytoskeleton, intermediate filaments and microtubules in smooth muscle cell migration. Respir Res. 18(1):54.

Theveneau E, Mayor R. 2013. Collective cell migration of epithelial and mesenchymal cells. Cell Mol Life Sci. 70(19):3481–3492.

Vasudevan HN, Soriano P. 2014. Srf regulates craniofacial development through selective recruitment of mrtf cofactors by pdgf signaling. Dev Cell. 31(3):332–344.

Walker BE, Fraser FC. 1956. Closure of the secondary palate in three strains of mice. Journal of Embryology and Experimental Morphology. 4(2):176–189.

Wilson NR, Olm-Shipman AJ, Acevedo DS, Palaniyandi K, Hall EG, Kosa E, Stumpff KM, Smith GJ, Pitstick L, Liao EC et al. 2016. Specc1l deficiency results in increased adherens junction stability and reduced cranial neural crest cell delamination. Sci Rep. 6:17735.

Xue G, Hemmings BA. 2013. Pkb/akt-dependent regulation of cell motility. J Natl Cancer Inst. 105(6):393–404.

Yu K, Ornitz DM. 2011. Histomorphological study of palatal shelf elevation during murine secondary palate formation. Dev Dyn. 240(7):1737–1744.

Zamir EA, Czirok A, Rongish BJ, Little CD. 2005. A digital image-based method for computational tissue fate mapping during early avian morphogenesis. Ann Biomed Eng. 33(6):854–865.

