## Supplemental Table and Figures for "SPECC1L-deficient palate mesenchyme cells show speed and directionality defect"

**Supplementary Table 1:** Significance values, established by Welch t-tests, of the differences among datasets shown in Figure 3n.

| **Parallel** | | | | |
|---|---|---|---|---|
| | wt low density | wt high density | mut low density | mut high density |
| wt low density | 0.500000 | 0.005978 | 0.028468 | 0.163239 |
| wt high density | 0.005978 | 0.500000 | 0.001777 | 0.006985 |
| mut low density | 0.028468 | 0.001777 | 0.500000 | 0.007666 |
| mut high density | 0.163239 | 0.006985 | 0.007666 | 0.500000 |

| **Perpendicular** | | | | |
|---|---|---|---|---|
| | wt low density | wt high density | mut low density | mut high density |
| wt low density | 0.500000 | 0.002246 | 0.386518 | 0.002792 |
| wt high density | 0.002246 | 0.500000 | 0.002170 | 0.014818 |
| mut low density | 0.386518 | 0.002170 | 0.500000 | 0.002689 |
| mut high density | 0.002792 | 0.014818 | 0.002689 | 0.500000 |

### Figure S1:

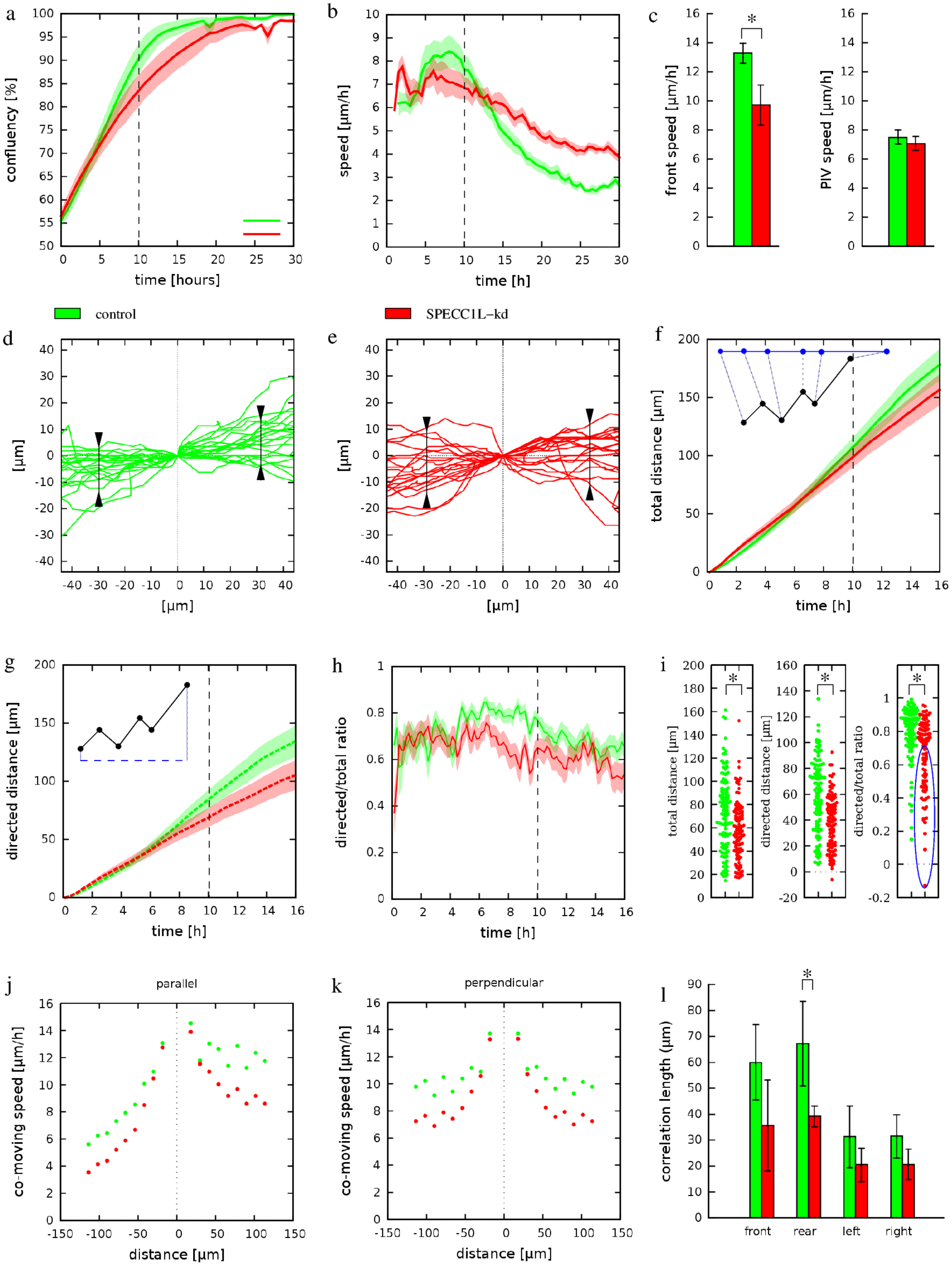
a
confluency [%]
time [hours]
b
speed [µm/h]
time [h]
c
front speed [µm/h]
PIV speed [µm/h]
control
SPECC1L–kd
d
e
[µm]
f
total distance [µm]
time [h]
g
directed distance [µm]
time [h]
h
directed/total ratio
time [h]
i
total distance [µm]
directed distance [µm]
directed/total ratio
j
parallel
co-moving speed [µm/h]
distance [µm]
k
perpendicular
co-moving speed [µm/h]
distance [µm]
l
correlation length (µm)
front
rear
left
right

**Figure S1: *SPECC1L*-kd U2OS cells show defects in directional migration. a)** Confluence analysis of wound-repair experiments indicated a delayed wound closure in *SPECC1L*-kd cell cultures (red) when compared to control cells (green). By 10h (dashed vertical line) control U2OS cells closed more than 75% of the wound. Solid lines represent the average of 17 independent microscopic fields, recorded in 4 distinct sets of experiments. Shaded areas indicate SEM. **b)** Average cell motility speed, calculated by PIV analysis of the 17 wound-closure recordings analyzed in panel (a). The motility of both control and *SPECC1L*-kd cells dropped as the wound closed. **c)** Statistical analysis of data presented in panels (a-b) indicated that while the wound-closure of *SPECC1L*-kd cells was significantly impaired in comparison to control U2OS cells ($p<5.5x10^{-6}$, Welch's t-test, left), there is no such profound difference in motile speed (right). Both the average speed of the wound-front and the average speed of cell movements were calculated within the 1h-15h time interval, prior to wound closure. Error bars represent SEM. **d-e)** Representative paths of individual cells moving near the wound-edge in control (d) and SPECC1L-kd (e) cultures. As in Fig. 2, trajectory heads were shifted to the origin, and 90% of the trajectories passed through the segments indicating the lateral spread of trajectories. Control cells exhibit a more directed movement to the open area compared to *SPECC1L*-kd cells. **f-h)** Statistical analysis of individual cell trajectories, in terms of total path-length (f), net directed displacement into the wound (g) and their ratio (h). Measures were plotted as a function of time elapsed, and indicate a reduction of both motile activity and directional guidance in *SPECC1L*-kd U2OS cells. Lines represent averages of 138 and 126 cells from control and *SPECC1L*-kd cultures, respectively. Shaded areas indicate SEM. Vertical dashed lines indicate the 10h timepoint, when the wound was almost closed. **i)** Statistical analysis of individual cell trajectories using the measures of total distance, directed displacement and their ratio, evaluated at 10h culture timepoint. The unguided population, delineated by blue ellipse, is more numerous in *SPECC1L*-kd cultures. The significance of differences was established by pairwise Welsh's t-tests, yielding p-values of $1.2x10^{-4}$, $2x10^{-9}$ and $1.1x10^{-7}$ for the total path length, directed displacement and ratio data sets, respectively. **j-k)** Collective movement of U2OS cells, indicated by average co-moving speed profiles (See Fig. 3) -- both along the front-rear (j) and lateral (k) axes. Data were averaged from 18 independent fields, both for control and *SPECC1L*-kd cells. Synchronized movement between *SPECC1L*-kd cells diminished at a closer distance than that of control cells. **l)** Average correlation lengths, both along directions parallel with and perpendicular to the direction of cell motion, were established by fitting an exponential function on the profiles shown in panels (j, k). Significance of the differences were established by Welch t-tests ($p<0.05$).

**Figure S2:**

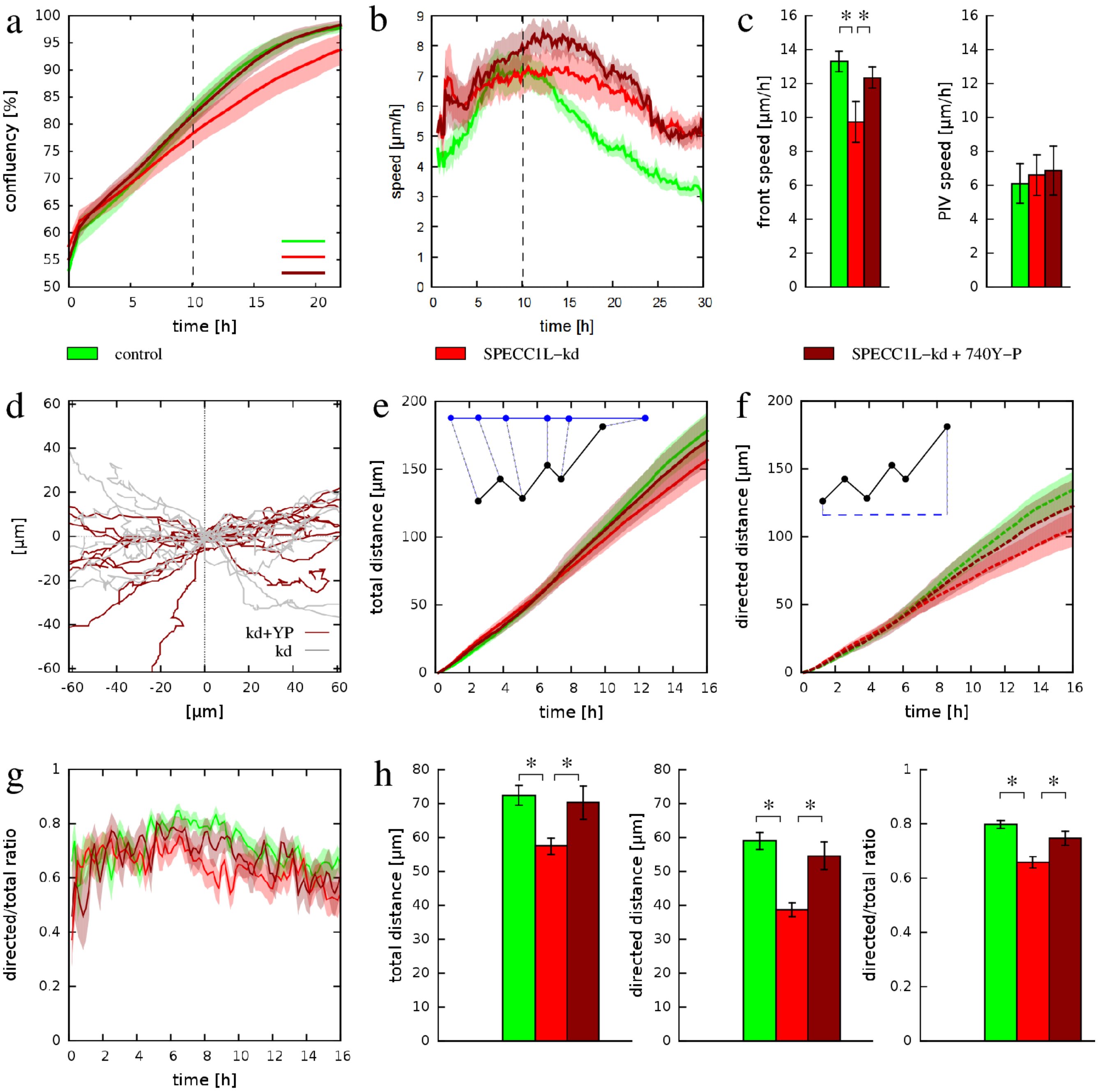
a
confluency [%]
time [h]
b
speed [µm/h]
time [h]
c
front speed [µm/h]
PIV speed [µm/h]
control
SPECC1L–kd
SPECC1L–kd + 740Y–P
d
[µm]
kd+YP
kd
e
total distance [µm]
time [h]
f
directed distance [µm]
time [h]
g
directed/total ratio
time [h]
h
total distance [µm]
directed distance [µm]
directed/total ratio

**Figure S2: Upregulation of PI3K-AKT signaling rescues both speed and directionality of *SPECC1L*-kd U2OS cells in wound-closure assays. a)** Confluence analysis of wound-repair experiments indicated that 100μg/ml 740Y-P enhances the wound-closure of *SPECC1L*-kd cells (dark red), compared to untreated *SPECC1L*-kd cultures (red). The improvement is enough to eliminate the difference with control cells (green). Solid lines represent the average of 17 independent microscopic fields, recorded in 4 distinct setd of experiments. Shaded areas indicate SEM. **b)** Average cell speed as a function of time, calculated from the 17 microscopic fields analyzed in panel (a). Solid lines represent averages, shaded areas indicate SEM. The PI3K-AKT activator elicited a sustained increase in the speed of *SPECC1L*-kd cells during the wound-closure process. **c)** Statistical analysis of wound-front propagation (left) and cell motility (right) speed data. Wound-front speed was obtained by linear fits over the confluency data within the 1h - 15h time-interval. Cell motility speed was obtained by averaging in the same time window. Error bars represent SEM. Significant differences are indicated by asterisks (control vs *SPECC1L*-kd: $p<5.5x10^{-6}$, 740Y-P treatment: $p<3.6x10^{-5}$). **d)** Representative trajectories of individual cells moving near the wound-edge in 740Y-P-treated *SPECC1L*-kd cultures (dark red). As a comparison, trajectories of untreated kd cells are plotted in grey. **e-g)** Statistical analysis of individual cell-trajectories, in terms of total path-length (e), net directed displacement into the wound (f) and their ratio (g). The PI3K-AKT activator improved each of these measures for *SPECC1L*-kd cultures (dark red vs red). Lines represent averages of 138, 126 and 73 cells from control, untreated *SPECC1L*-kd cultures, and 740Y-P-treated *SPECC1L*-kd cultures, respectively. Shaded areas indicate SEM. **h)** Statistical analysis of individual cell-trajectories using the measures of total path-length, directed displacement and their ratio, evaluated at 10h culture timepoint. The significance of 740Y-P treatment was established by pairwise Welsh's t-tests, yielding p-values of 0.02, $6.6x10^{-4}$ and 0.01 for the total path length, directed displacement and ratio data sets, respectively. Error bars indicate SEM.
